# De Bruijn Graph Partitioning for Scalable and Accurate DNA Storage Processing

**DOI:** 10.1101/2025.05.19.654814

**Authors:** Florestan De Moor, Olivier Boullé, Dominique Lavenier

## Abstract

**Motivation:** DNA-based data storage offers a compelling solution for long-term, high-density archiving. In this framework, accurately reconstructing high-quality encoded sequences after sequencing is critical, as it has a direct impact on the design of error-correcting codes optimized for DNA storage. Furthermore, efficient and scalable processing is essential to manage the large volumes of data expected in such applications.

**Results:** We introduce a novel method based on de Bruijn graph partitioning, enabling fast and accurate processing of sequencing data regardless of the underlying sequencing technology and without requiring prior knowledge of the information encoded in the oligonucleotides. Evaluated on both synthetic and real datasets, the method achieves excellent precision and recall. It is implemented in C++ within the software ConCluD and optimized for multi-core servers. Our experiments demonstrated that a dataset of 55 million reads, corresponding to a 135 MB binary file, can be processed in less than 10 minutes on a 16 hyper-threaded core server.

## 1. Introduction

The world is experiencing an exponential surge in digital information, posing significant challenges for archival storage. Existing technologies consume large amounts of energy, occupy substantial physical space, incur high infrastructure costs, and require extensive maintenance to ensure long-term data preservation. Hence, new storage solutions need to be found. Advances in molecular synthesis and the rise of efficient sequencing technologies have opened the possibility of storing data in DNA molecules. This approach is gaining increasing interest as a potential solution to overcome the current technological barriers hindering its widespread adoption [1] [2] [3] [7].

DNA-based data storage follows several key steps. First, binary files are encoded into a DNA sequence representation. Then, DNA molecules are synthesized based on this encoded data and stored in a protected environment. To retrieve the information, the process begins by selecting the DNA molecules that correspond to the desired files. These molecules are then sequenced, and the resulting data are transmitted to the decoding stage to reconstruct the original files.

Current synthesis processes can only produce short oligonucleotides with a maximum length of about 300 nucleotides. As a result, files must be divided into smaller fragments, each of which is indexed to allow the reconstruction of the original information. For example, a video file would be fragmented into hundreds of millions of oligonucleotides.

In a single storage unit, thousands of different documents can co-exist. To identify them, each file can be assigned a unique nucleotide sequence as a tag. This tag not only helps distinguish files, but can also be used to biochemically select the oligonucleotides associated with a specific file. In this case, the tag can be designed as a pair of primers, which enables the targeted amplification of the molecules using PCR.

In this framework, reading a file involves selecting the corresponding molecules by performing a PCR using the specific pair of primers associated with it, followed by sequencing the PCR product.

However, the sequencing output cannot be sent directly to the decoding stage. The data must first be processed to eliminate redundancy introduced during sequencing and to reconstruct the initial set of sequences that represent the file as accurately as possible. The first processing step involves clustering the reads to group similar sequences together. In the second step, a consensus sequence is established within each group, allowing for the correction of potential errors. This process ensures the production of high-quality sequences, which can then be transmitted to the decoder stage.

Since sequencing is a stochastic process, it is highly likely that not all sequences will be successfully reconstructed, and some of those provided to the decoder may still contain errors. To address this issue, the encoding process incorporates error-correcting codes. These codes enable the reconstruction of the complete information from only a subset of recovered sequences while also correcting residual errors, ensuring data integrity [4] [5] [6].

The work presented here focuses on processing sequencing data with the goal of delivering as many correct sequences as possible to the decoder. Similar work for genomic purpose has already been carried out, mainly in bioinformatics and metagenomics, but they mainly focuses on DNA clustering algorithms. Methods such as UCLUST [8], CD-HIT [9] and DNACLUST [10] use greedy algorithms for sequence similarity, while MeShClust [11] and MetaDEC [12] use machine learning approaches. Graph-based techniques, such as MMseqs2 [13], offer further strategies. However, a common limitation of these algorithms is their reliance on a fixed sequence identity threshold, which can be difficult to determine, particularly in DNA storage systems. Unlike genomic clustering, DNA data storage presents unique constraints due to shorter DNA strands (100-300 nucleotides). Although algorithms such as ALFATClust [14] and SEED [15] attempt to dynamically adjust clustering parameters, they remain optimized for metagenomic data. Recent advances in large-scale DNA clustering for storage applications include the work of Rashtchian et al. [16] and GradHC [17], both of which are capable of clustering billions of reads and producing highly reliable clustering results. The output of these programs are clusters containing a small number of reads, which are then used to build a consensus sequence.

Our approach is distinct and aims to merge the clustering and consensus steps into a single process. To achieve this, we draw inspiration from bioinformatics techniques used in genome assembly, particularly the use of de Bruijn graphs [18] [19]. This method has already been explored by [20] and has proven to be particularly robust in handling DNA breaks, rearrangements, and indels. However, as the size of the encoded files increases, the construction of large de Bruijn graphs can lead to excessive computation times.

Our method reduces algorithmic complexity by hierarchically dividing the initial set of reads into smaller subsets of predefined size. Each subset is expected to contain a portion of the original sequences along with their noisy copies. A de Bruijn graph is then constructed for each subset. Given the known length (N) of the target sequences and the nucleotide sequences that mark their start and end (the primers), the sequences to be reconstructed correspond to all paths of length N between two k-mers derived from the start and end primers. The advantages of this method are: (a) the hierarchical partitioning is extremely fast; (b) constructing smaller de Bruijn graphs I (c) parallelization on multicore architectures is straightforward since there are no dependencies between the de Bruijn graphs.

## 2 Method

In the context of DNA storage, reference sequences consist of three concatenated parts, as illustrated in Figure 1: a start primer, a payload, and an end primer. The payload carries the encoded data, along with indexing information for document reconstruction, optional checksums, and error correction data. To retrieve a document, a PCR is first performed using the specific primer pair associated with it, followed by a sequencing process. The resulting dataset may contain billions of reads. Here, we assume that each read includes a complete sequence, encompassing both primers.

**Figure 1:**
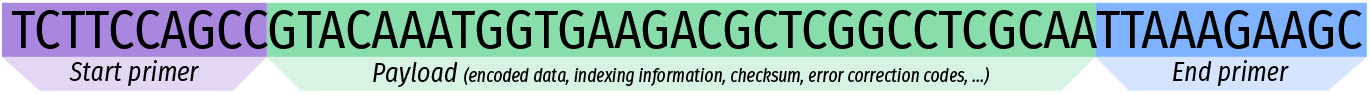
Structure of a reference sequence in the context of DNA storage

To recover the initial set of sequences, a straightforward strategy would be to construct a de Bruijn graph using all these sequences. Since all sequences share the same start and end primers, one could then enumerate all paths in the graph that begin with a k-mer extracted from the start primer and end with a k-mer from the end primer. Each path represents a possible sequence. Paths that do not have the correct length are discarded. In practice, this solution works well and remains relatively fast for graphs of reasonable size. However, as the number of sequences increases, the computation time grows proportionally. Indeed, constructing a de Bruijn graph involves k-mer counting, which is a computationally intensive task. Also, identical k-mers in different sequences lead to an increase in the number of possible paths, drastically raising the complexity of the problem.

To mitigate this complexity, our method first partitions the dataset (fastq file) into multiple subsets of approximately equal size. Each subset, and it is of the most importance, contains all instances of a given original sequence (see Section Partitioning). From there, as mentioned earlier, the problem consists of searching for paths in graphs that are now of reduced size.

The algorithm represented figure 2 can be abstracted as follows:

**Figure 2:**
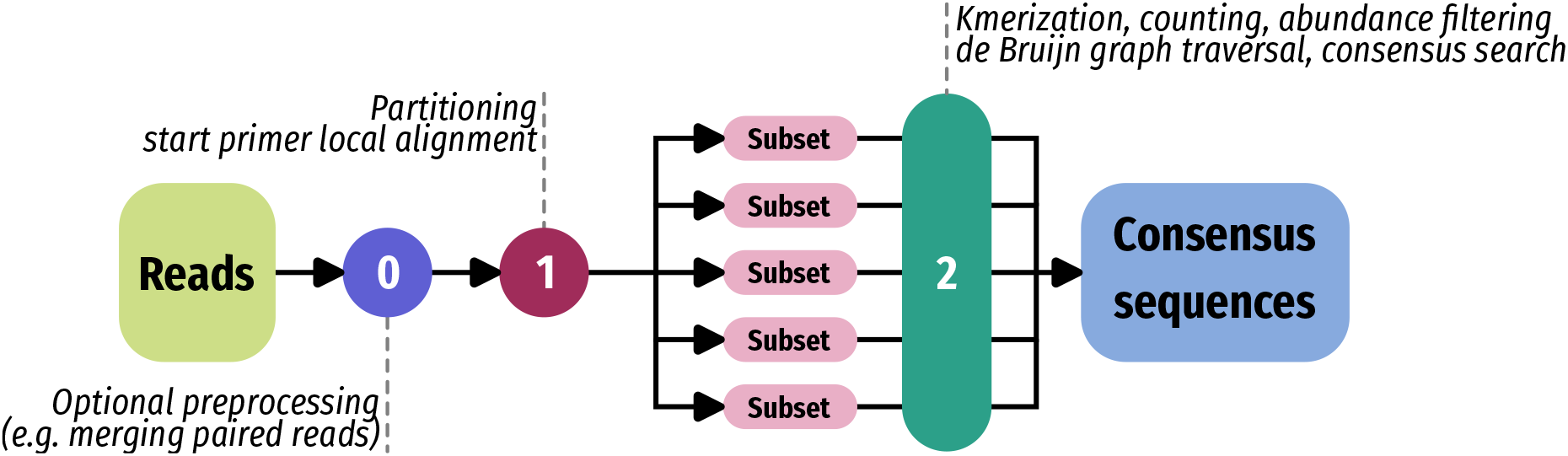
Overview of our method. Step 1 aligns the start primers and partition all reads. Step 2 builds de Bruijn graphs. Step 3 traverses the graphs to find consensus sequences of the expected length.

1. Partition the dataset into subsets of approximate equal size
2. For each subset
  - Create a de Bruijn graph
  - Generate consensus sequences from the graph

To accelerate computation in a multithreaded environment, step 1 can benefit from intra-read parallelism, while step 2 allows for intra-subset parallelism. The following subections decribe in detail each of these steps.

### 2.1 Partitioning

The objective of this step is to distribute the entire set of reads into subsets so that all copies of the same sequence are present in only one subset. The first task is to identify the start primer within the read, considering that the method is independent of the sequencing technology and that the primer is not necessarily located at the very beginning of the read. To achieve this, a local alignment is performed for each read against the start primer (and its reverse complement). However, some reads may be lost if no sufficiently good alignment is found. This is more likely to occur for reads with a high error rate. But removing them is actually beneficial, as they could otherwise compromise the accuracy of the consensus search.

From the primer alignment position, a few bases immediately following the primer are used as a partitioning key. To account for the unknown data distribution, we use a dynamic key size. This approach ensures a more balanced load distribution and prevents some subsets from having an excessive number of reads while others remain nearly empty. We start with keys of 4 bases and then recursively split each subset into 64, assigning one subset to each possible value of the next 3 bases, thereby expanding the key. This process continues until each subset contains no more than a predefined number of reads. Ultimately, this step produces a set of FASTA files, each containing approximately the same number of reads.

Each subset is written to disk and reads are trimmed to eliminate irrelevant parts: segments preceding the starting primer and segments after the expected length of the sequence are removed. This trimming can be done because the characteristics of the original reference are known, in particular the length of the oligonucleotides that encode a file.

More generally, the partitioning strategy accept several start primers as large files are often indexed in several parts, each identified by a specific pair of primers. Partitioning is therefore initially based on these different start primers.

This partitioning method is very fast and efficient. It systematically groups reads with similar start sequences. With a low error rate, the majority of copies of the same sequence will end up in the same subset. However, errors on the first bases of the read will inevitably lead to incorrect read assignment. But with current error rates and advances in sequencing technologies and base-calling algorithms, the number of misassigned reads remains fairly low, and does not disrupt processing in subsequent steps.

### 2.2 de Bruijn graph building

De Bruijn graphs are primarily built based on k-mer counting. The advantage of considering only small subsets (fewer than 10,000 reads) is that the resulting graph will also be of limited size. The k-mer length can be optimally adjusted, enabling extremely simple and fast counting techniques that do not require complex data structures. In this case, a simple hash table is used and the values of the counters are capped to 255 to fit into 8-bit variables.

Finally, we discard the k-mers that appear less than a user-defined threshold. This filtering eliminates sequencing errors, but also k-mers from reads assigned to an improper subset.

### 2.3 Generating consensus sequences

The last stage consists of considering the k-mers previously found as a de Bruijn graph and traversing its paths to find consensus sequences.

The exploration begins at the node representing the starting k-mer, which is actually the rightmost k-mer of the start primer. Then, the next possible four bases are considered recursively. If the traversal reaches the leftmost k-mer of the ending primer and the current payload has the exact expected size, the sequence is considered a consensus. If the traversal results in a sequence that exceeds the expected consensus length, the search on that particular path is terminated.

Since this exploration has exponential complexity, due to cycles in the graph, several techniques are employed to reduce processing time. First, the graph is progressively pruned by removing nodes with no outgoing edges. Second, when a path is terminated, the current sequence is analyzed for repeated k-mers. If repetitions are detected, one edge is removed from the graph to break the loop. This optimization is lossy, meaning that it may prevent some valid consensus sequences from being found. However, eliminating loops significantly accelerates the search process. As in any data transmission context, some sequences are expected to be missing. To address this, efficient techniques such as error correction codes can be applied.

## 3 Evaluation setup

### 3.1 Hardware

Hardware-wise, we use a server fitted with an Intel® Xeon® Silver 4215 CPU @ 2.5 processor (Skylake architecture, 16 hyper-threaded cores) and 256 of DDR4 @ 2.4 RAM. The server operates on Debian 10.

### 3.2 Software

**ConCluD** (<u>C</u>onsensus algorithm based on <u>Clu</u>stering for DNA storage <u>D</u>ecoding) is the name of the C++ software developed for the method described in the previous section. Unless state otherwise, ConCluD is executed with 32 CPU threads with the following default parameter values:

- k-mer size = 15
- k-mer abundance threshold = 3
- maximum number of reads in a subset = 5000

**DBGPS** has been developed by [20] and also uses a de Bruijn graph approach to reconstruct sequences after sequencing. A single de Bruijn graph is built from the whole read dataset. To reconstruct sequences, the oligonucleotides must be necessarily structured into three distinct fields: an index area, followed by the payload, and then a CRC field. The entire sequence is also framed by two primers. The C multithreaded version is used for comparison.

**GradeHC**: has been developed by [17] and only generates clusters of reads. Each cluster can then be used to generate a single consensus sequence.

### 3.3 Datasets

This section presents the different datasets we used for evaluating ConCluD. The following table summarizes the synthetic datasets. For each dataset, it gives its name, the size of the oligonucleotides, the sequencing technology, the number of primers, the number of oligonucleotides and the resulting reads with an average coverage of 30x. A detailed description of each of them, and the way they have been designed, can be found in annex.

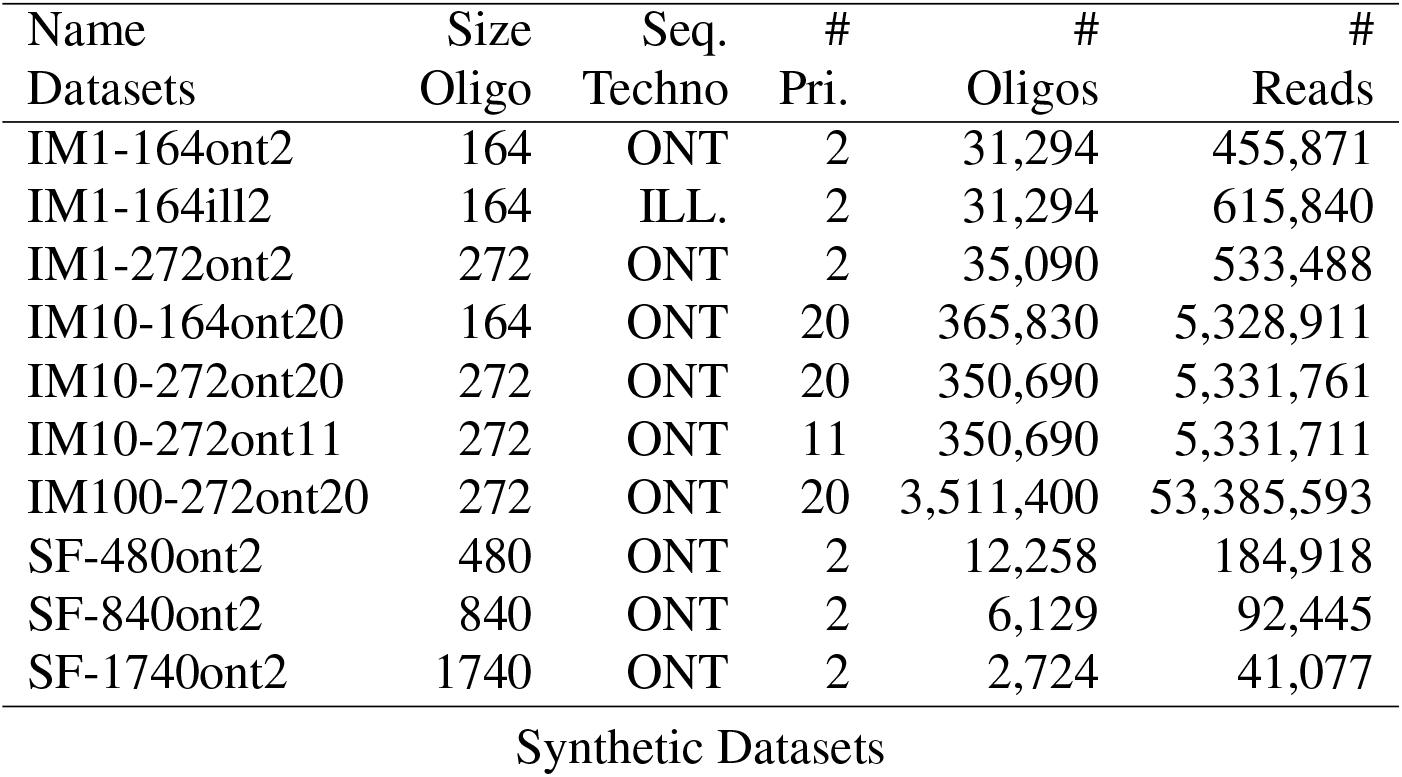

| Name<br>Datasets | Size<br>Oligo | Seq.<br>Techno | #<br>Pri. | #<br>Oligos | #<br>Reads |
| --- | --- | --- | --- | --- | --- |
| IM1-164ont2 | 164 | ONT | 2 | 31,294 | 455,871 |
| IM1-164ill2 | 164 | ILL. | 2 | 31,294 | 615,840 |
| IM1-272ont2 | 272 | ONT | 2 | 35,090 | 533,488 |
| IM10-164ont20 | 164 | ONT | 20 | 365,830 | 5,328,911 |
| IM10-272ont20 | 272 | ONT | 20 | 350,690 | 5,331,761 |
| IM10-272ont11 | 272 | ONT | 11 | 350,690 | 5,331,711 |
| IM100-272ont20 | 272 | ONT | 20 | 3,511,400 | 53,385,593 |
| SF-480ont2 | 480 | ONT | 2 | 12,258 | 184,918 |
| SF-840ont2 | 840 | ONT | 2 | 6,129 | 92,445 |
| SF-1740ont2 | 1740 | ONT | 2 | 2,724 | 41,077 |

Synthetic Datasets

The data are from high-resolution images. The payloads were constructed in compliance with the requirements for DNA storage (no large homopolymers, GC content around 50%, primers that do not hybridize in the middle of the payload, etc.). In addition, to ensure compatibility with the DBGPS dataset format, the same oligonucleotide structure was used for the IM1, IM10 and IM100 datasets, although ConCluD does not need any indexes to process the data. The ONT and Illumina sequencing datasets have been respectively generated with the PBSIM3 [22] and ART simulator tools [21]. The coverage has been approximately set to 30X.

The three datasets IM1-xxx have been designed from small size JPEG images (497 KB and 1.3 MB) encoded with oligonucleotides of sizes 164nt and 272nt and synthetized with ONT or Illumina read simulators. Each image is encoded within a single pool of oligonucleotides sharing the same start and end primers, leading to 2 primers to perform a PCR.

The 3 datasets IM10-xxx have been designed from larger images (5.8 MB and 13.5 MB) encoded with oligonucleotides of size 164 nt and 272 nt. Each image is encoded within 10 pools of oligonucleotides with 2 variants: for IM10-xxx20 each pool have a different pair of primers, leading to a total of 20 primers (10 *×* 2). For IM10-ont11, each pool have the same start primer, but a different end primer, reducing the number of primers to 11.

The dataset IM100-272ont20 has been designed from a single very large JPEG image (134.7 MB) which is encoded into 100 different pools of oligonucleotides of size 272 nt. Each pool is distinguished by a different pair of primers among 10 start and 10 end primers.

The reason for splitting the binary file into small chunks of oligonucleotide pools is that it will make the decoding process simpler and faster in the downstream steps. Running error correcting codes on small pools, especially to retrieve lost oligonucleotides, is much less complex than doing it on large pools. In this experimentation it is also a way to investigate the best choice regarding pair of primers. We will see in the next section that sharing primers provides better performances.

The datasets SF-xxx refer to the same image, but encoded with oligonucleotides of size 480nt, 840nt and 1740nt. The purpose of these three datasets is to test how ConCluD performs on large oligonucleotides. Since DBGPS primarily targets small oligonucleotides, these datasets won’t be used for comparison.

To test and evaluate ConCluD on real data, the following real datasets have been selected:

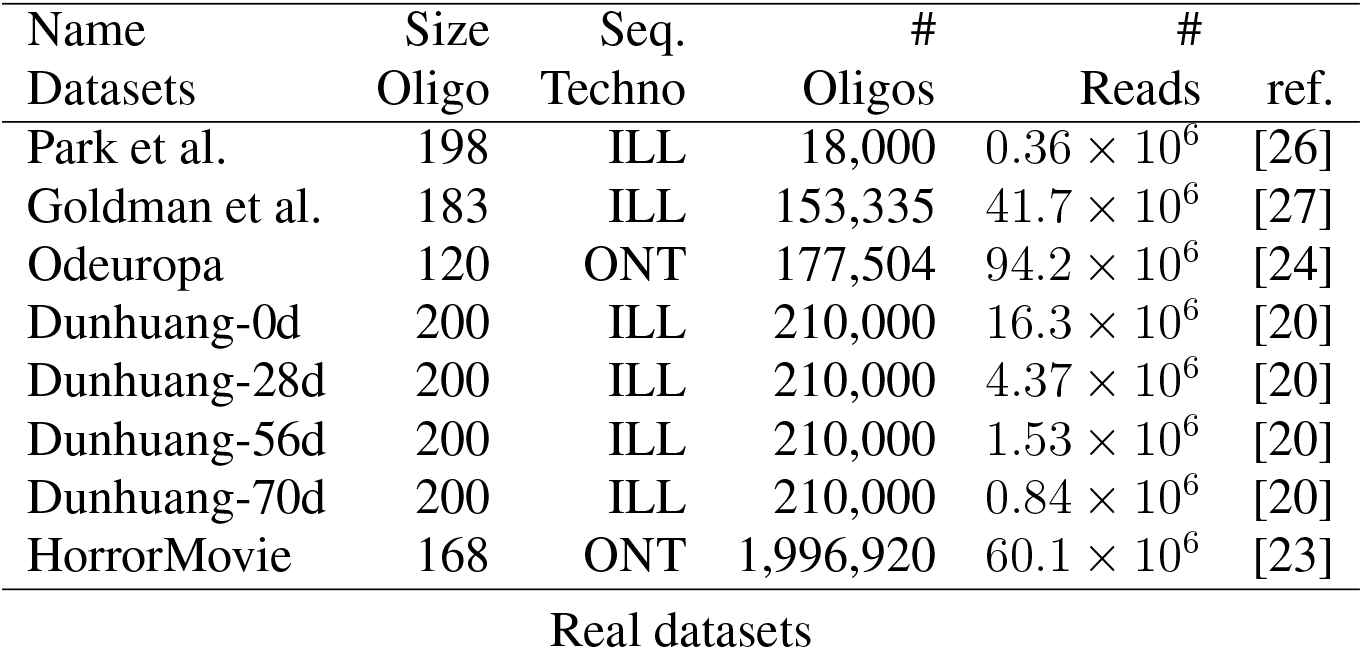

| Name | Size | Seq. | # | # |  |
| --- | --- | --- | --- | --- | --- |
| Datasets | Oligo | Techno | Oligos | Reads | ref. |
| Park et al. | 198 | ILL | 18,000 | $0.36 \times 10^6$ | [26] |
| Goldman et al. | 183 | ILL | 153,335 | $41.7 \times 10^6$ | [27] |
| Odeuropa | 120 | ONT | 177,504 | $94.2 \times 10^6$ | [24] |
| Dunhuang-0d | 200 | ILL | 210,000 | $16.3 \times 10^6$ | [20] |
| Dunhuang-28d | 200 | ILL | 210,000 | $4.37 \times 10^6$ | [20] |
| Dunhuang-56d | 200 | ILL | 210,000 | $1.53 \times 10^6$ | [20] |
| Dunhuang-70d | 200 | ILL | 210,000 | $0.84 \times 10^6$ | [20] |
| HorrorMovie | 168 | ONT | 1,996,920 | $60.1 \times 10^6$ | [23] |

Real datasets

The Park dataset was designed using Fountain codes to explore how low-quality reads can be leveraged to reduce sequencing coverage requirements [26]. The Goldman dataset [27] includes five files totaling 757 KB of data, in which text, images, and audio have been encoded. The Odeuropa dataset is provided by the OligoArchive European project [24], and the Dunhuang-dX datasets originate from [20] where ten digital pictures of Dunhuang murals have been encoded. Each Dunhuang datasets is based on the same initial sample but was subjected to accelerated aging for 0, 28, 56, or 70 days to induce molecular degradation. Finally, the HorrorMovie dataset is derived from Malcolm Le Grice’s Horror Film 1, which was encoded into nearly 2 million of nucleotides [23].

Illumina sequencing generates paired-end reads, which need to be merged to reconstruct complete oligonucleotides. This step is crucial for our strategy, as it requires both the start and end primers to be present on the same read. Merging was performed using the NGmerge tool [25]. However, in some datasets, primer or adapter sequences were missing due to trimming during the base calling process, which fortunately reorients all sequences identically. To maintain compatibility with our approach, artificial start and end primers have been added to restore the expected read structure.

### 3.4 Metrics

To evaluate the accuracy of our algorithm, we consider the precision and recall metrics.

Precision measures the amount of sequences returned by the program that are actually present in the reference. Achieving a good accuracy is important to ensure the reconstruction does not alter the original data. We compute the raw precision of the algorithm, independently of any checksum that might be embedded into the payload, and the final precision that allows to discard sequences after checksum verification.

On the other hand, recall measures the proportion of the reference that was found by the program. A bad recall hurts the reconstruction of information as some data blocks may be missing. We compute the raw recall of the program, independently of any error correction codes or other device that might be embedded into the payload to recover missing sequences.

Concerning time performance, we report the elapsed time of any execution.

## 4 Evaluation results

### 4.1 Precision and Recall

Figure 3 reports the precision and recall of ConCluD and DBGPS software for each datasets. ConCluD parameters (k-mer size, k-mer threshold, subset size), have been set to optimized the results according to the dataset features. Overall, on small oligonucleotides (164nt and 272nt), ConCluD achieves excellent results in both precision and recall. On the other hand, DBGPS only produces satisfactory results for small datasets with limited oligonucleotide sizes. For the five last datasets, DBGPS does not output any consensus. More specifically, for the SF datasets, DBGPS crashes with a stack smashing error. For dataset IM10-272ont11, the execution time is excessively long (24 minutes) and the output consensus sequences are erroneous. And for dataset IM100-272ont20, the process was killed due to memory overflow (256 GB). Indeed, in DBGPS, the size of the de Bruijn graph is directly proportional to the number of reads (and thus to the number of k-mers). The complexity of traversing the graph and reconstructing paths increases accordingly. Removing loops in the graph to reduce complexity has an immediate impact: many paths supporting reference sequences are not explored. ConCluD integrates the same strategy but applies it to a multitude of small graphs, where only a few loops are removed, leading to marginal impact on the results.

**Figure 3:**
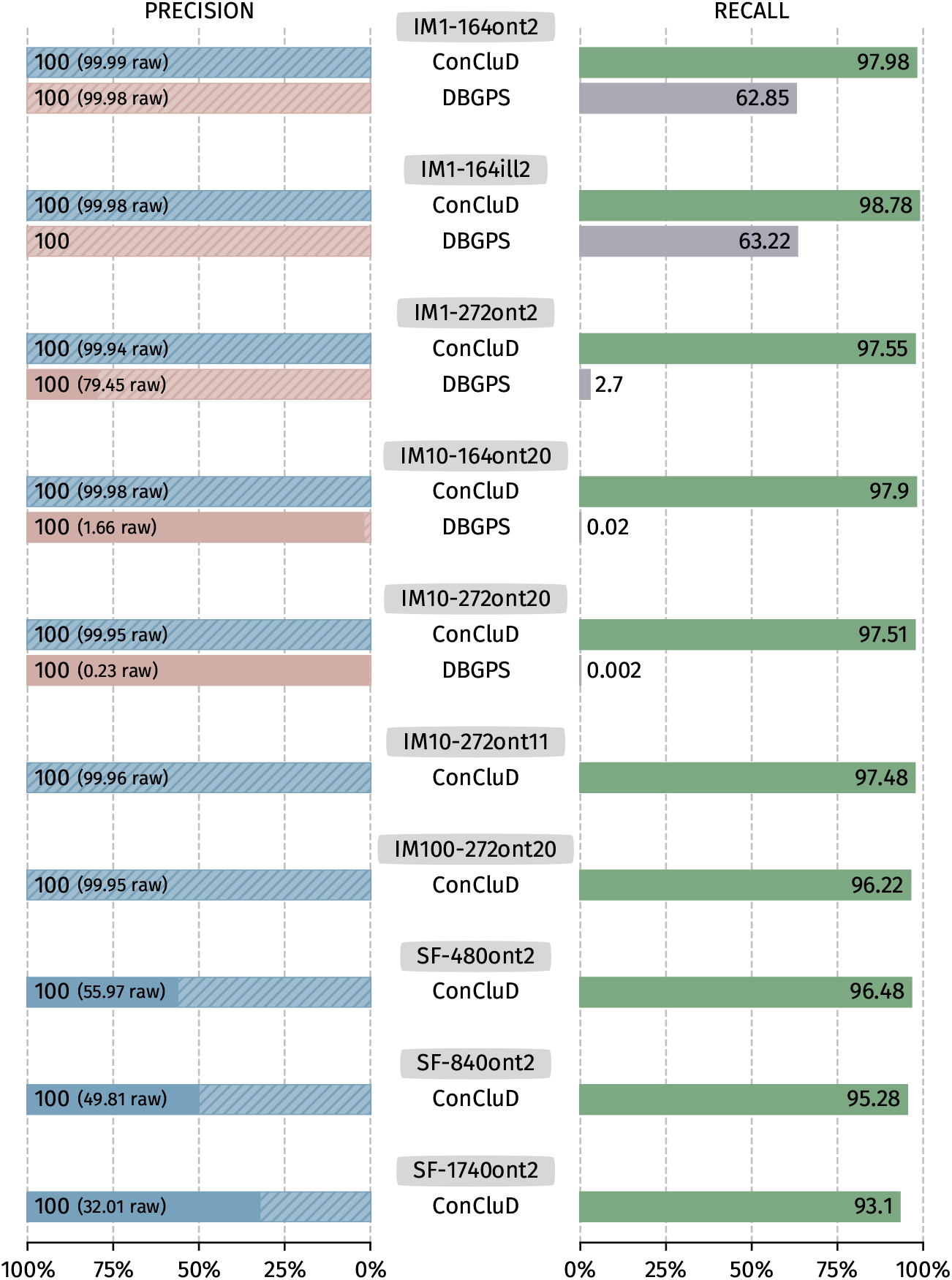
Precision and Recall on the synthetic datasets. Raw precision measures the amount of sequences returned by the software without CRC. Overall precision, actually 100% on each datasets, take care of CRC

As previously mentioned, the oligonucleotides were structured to be compatible with the DBGPS format. ConCluD relies on the nucleotide sequence directly following the starting primer, which, in this case, corresponds to the added index and could introduce bias into our strategy. Therefore, we tested our partitioning technique by using the partitioning position after the index, which corresponds to the beginning of the payload. The results remain identical. No bias is introduced.

The last three datasets (SF-xxx) test the robustness of ConCluD against long oligonucleotides. A slight decrease in recall is observed when the length of the oligonucleotides increases. On the raw precision side, however, a very clear decrease is observed, which is consistently compensated by a validity check.

This evaluation shows that achieving 100% recall is never possible, either because on large oligo pools the synthesis did not produce all the expected oligonucleotides or because the random sequencing process failed to capture all the information. This is why error-correcting codes are added to recover the information. Although this is not the focus of this work, these results may help quantifying sequencing losses and adjusting the parameters of error-correcting codes.

### 4.2 Execution time

Figure 4 shows the execution times for ConCluD and DBGPS. For the datasets IM10-272ont11 and IM100-272ont20, DBGPS execution times are not reported as no results are produced. For ConCluD, two things can be observed:

**Figure 4:**
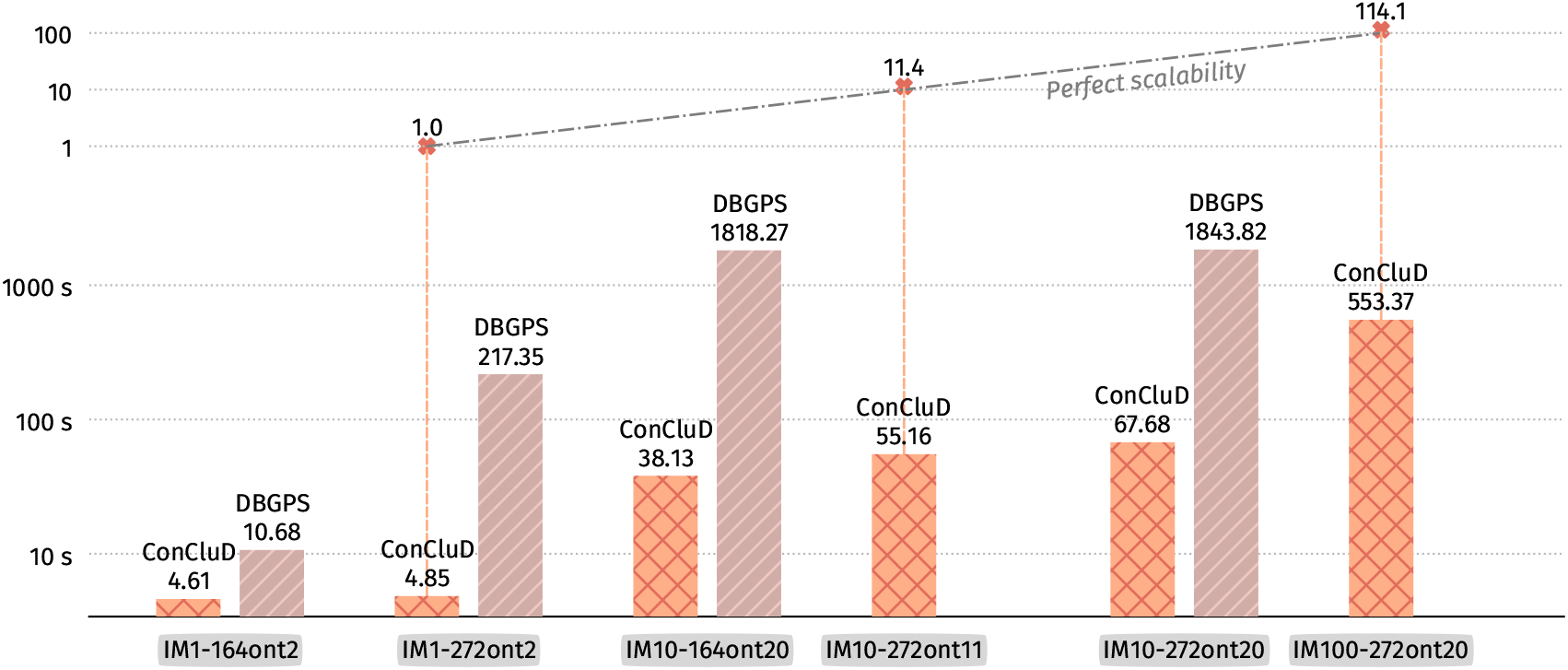
Execution time for ConCluD and DBGPS.

1. The execution time increases almost linearly with the data size. The top of the figure shows the relative execution time using the smallest dataset execution time as reference.
2. The difference between IM10-272ont11 (with a common start primer for all images) and IM10-272ont22 (with different start and end primers for each image) favors the dataset using only 11 primers. There is a significant decrease, which can be explained by the optimization of the graph traversal. Instead of performing 10 searches with 10 different start primers, only one search is initiated. What’s more, instead of performing 10 alignments for each primer, we do just one, so partitioning also involves 10x fewer calculations. Moreover, it’s better to have 1 start primer and 10 end primers, than 10 start primers and 1 end primer. Behavior is not symmetrical.

GradHC, that only performs clustering for DNA storage, has been run on dataset IM1-164ont2 (1 image - 31,294 oligonucleotides). The measured execution time was 2580 seconds. This time is actually far too long for this method to be practically used.

ConCluD has two major steps: (1) partitioning the dataset; (2) building consensus sequences from a de Bruijn graph. The table below indicates the execution time (in second) for each step:

|  | IM1<br>272ont2 |  | IM10<br>272ont11 |  | IM100<br>272ont20 |  |
| --- | --- | --- | --- | --- | --- | --- |
| Partitioning | 4.52 | (93.2%) | 41.17 | (74.6%) | 486.94 | (88.0%) |
| Consensus | 0.33 | (6.8%) | 13.99 | (25.4%) | 66.43 | (22.0%) |
| Total | 4.85 |  | 55.16 |  | 553.37 |  |

Distribution of execution times, in second, between the two phases of ConCluD. The values in brackets indicate the percentage of time spent in the 2 stages.

We immediately notice that the partitioning step is dominant. The construction of de Bruijn graphs (mainly k-mer counting) is fast, as well as the path search within these different graphs.

### 4.3 Parameter impacts

There are three main parameters that govern the operation of ConCluD: the size of the subsets in the partitioning phase, the size of the k-mers and their solidity in the consensus phase. These parameters can influence the quality of the results (precision and recall) as well as the execution time. Experimental tests show that a subset size of 5,000 reads, a k-mer size of 15 nt, and a low-abundance k-mer threshold of 3 provide good results and can be used as default parameters. In the following sections, the impact of these parameters is evaluated separately, with the others set to their default values.

Figure 5 evaluates the impact of subset size using three different values: 5K, 25K, and 50K reads. The results clearly show that larger subsets significantly reduce recall. This supports our strategy of managing multiple small de Bruijn graphs rather than a single large one. The execution time is less affected; partitioning into larger batches is slightly faster. However, the processing time for consensus finding increases significantly. A maximum subset size of 5K reads therefore appears to be an optimal choice.

**Figure 5:**
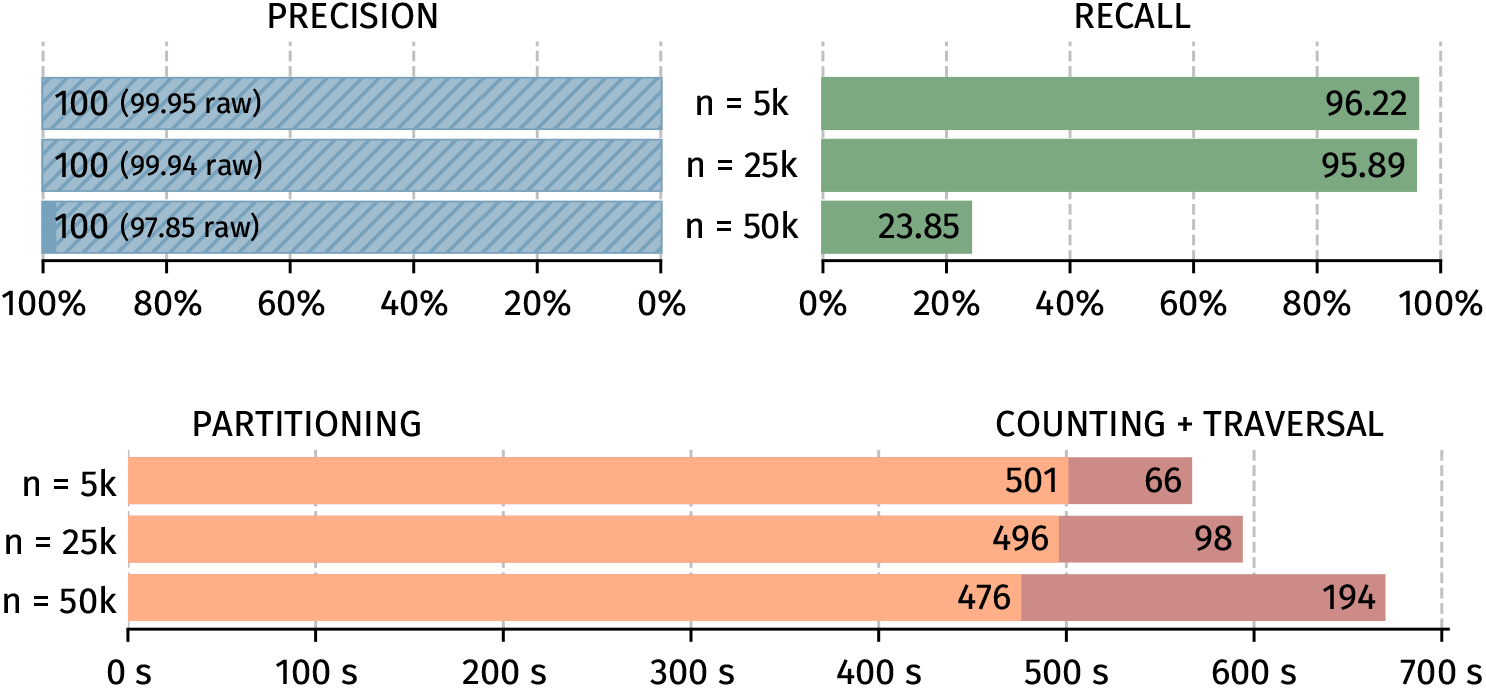
Impact of the subset size for precision, recall and execution time

The size of the k-mers (figure 6) have a strong impact. As it can be seen the minimum size is 13nt. Above this value, both precision and recall are good, and the computation time remains stable. A default value of 15 is therefore entirely appropriate.

**Figure 6:**
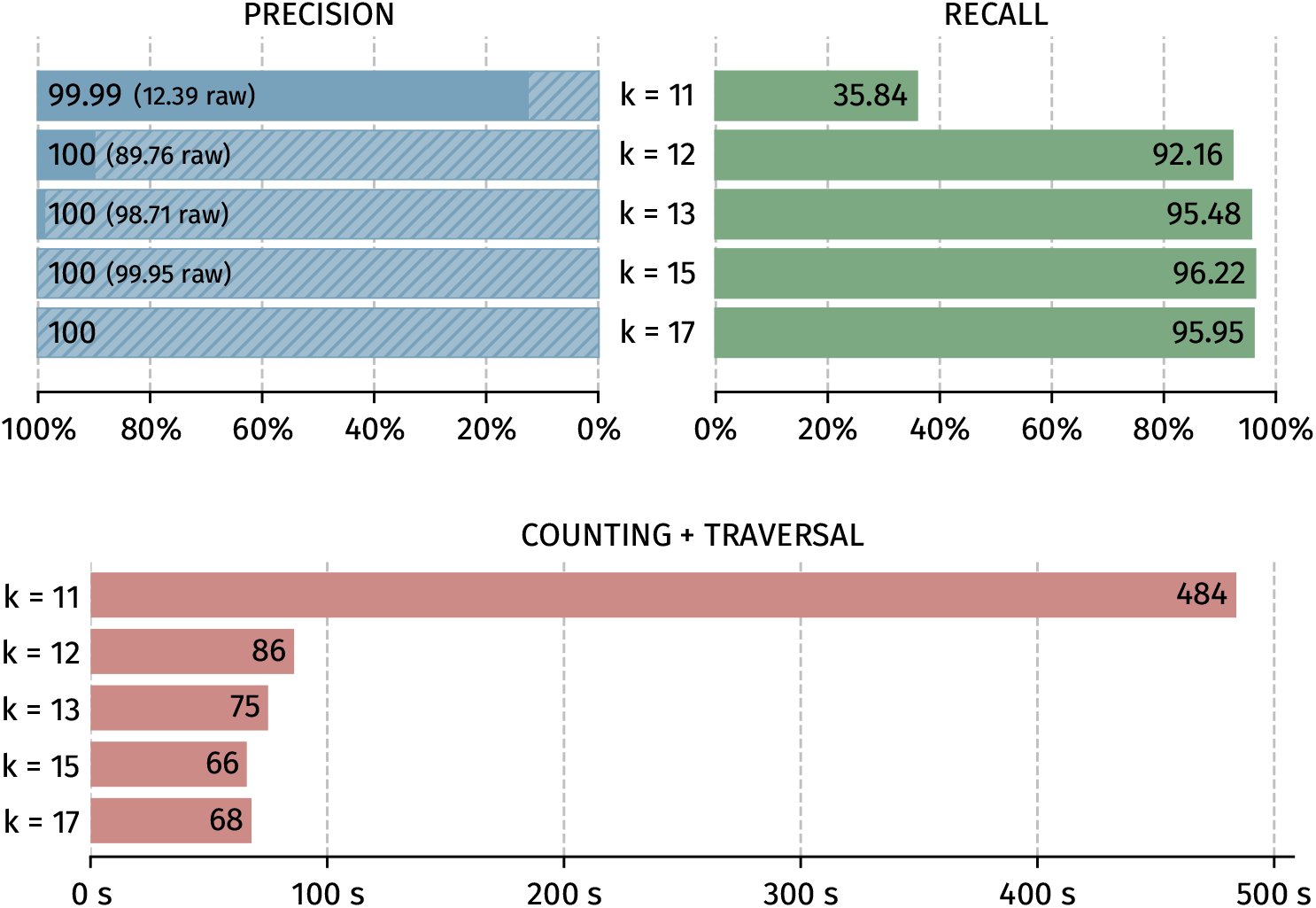
Impact of the k-mer size for precision, recall and consensus execution time

If the value of the low-abundance k-mer threshold (Figure 7) used to build the de Bruijn graph is too low (i.e. set to 1) too many false positive sequences are generated, leading to very poor precision and excessive computation time due to very noisy de Bruijn graph. On the other hand, as this value increases, the recall decreases accordingly. A good balance to optimize the recall is achieved with a trade-off value of 3 for an expected coverage of 30X.

**Figure 7:**
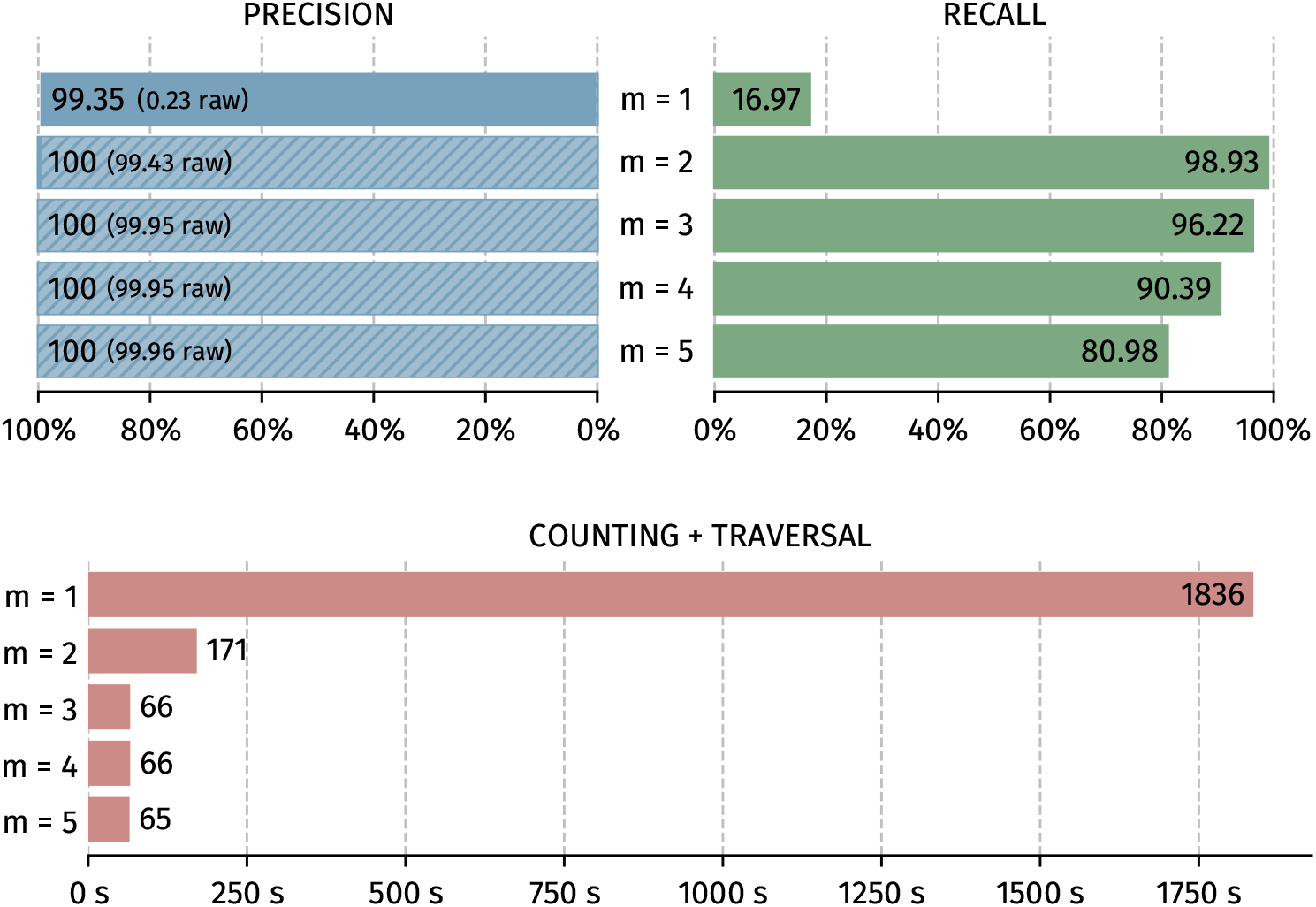
Impact of the k-mer abundance threshold for precision, recall and consensus execution time

### 4.4 Evaluation on real datasets

Finally, ConCluD was run and evaluated on real datasets, which were subsampled when required to ensure a maximum coverage of 30X. The results are presented in the following table.

|  | % Precision | % Recall | Exec. time |
| --- | --- | --- | --- |
| Park et al. | 99.87 | 99.99 | 0.77 sec. |
| Goldman et al. | 70.66 | 97.21 | 14.61 sec. |
| Odeuropa | 96.52 | 99.47 | 8.78 sec. |
| Dunhuang-d0 | 91.26 | 98.73 | 40.90 sec. |
| Dunhuang-d28 | 91.24 | 81.41 | 13.69 sec. |
| Dunhuang-d56 | 86.63 | 19.94 | 5.54 sec. |
| Dunhuang-d70 | 86.86 | 7.89 | 2.92 sec. |
| HorrorMovie | 99.99 | 94.00 | 145.00 sec. |

For the Dunhuang-dX datasets, we report only the raw precision, as the verification protocol is unavailable. However, when this information is accessible, incorporating it typically increases the precision to 100%. Recall remains high, except for degraded sequences in the Dunhuang datasets. In such cases, our strategy becomes ineffective, as the molecular damage fragments the sequences, preventing ConCluD from performing consistent partitioning. The experiments conducted by the authors of DBGPS represent extreme conditions, and it is reasonable to assume that a well-designed archival system would avoid such scenarios.

## 5 Conclusion

We presented a method based on de Bruijn graphs to process sequencing reads derived from oligonucleotides and reconstruct the DNA sequences encoding a digital document. The method has been implemented in the ConCluD software.

The originality and efficiency of our approach lie in a partitioning strategy that groups each sequence and all of its copies into the same subset. Each subset is then processed independently. This offers two key advantages: first, the resulting graphs are smaller, which simplifies their construction and handling; second, this execution model allows for highly scalable processing. Our strategy is also compatible with longer oligonucleotide lengths, anticipating future technological advances that could enable the storage and retrieval of significantly larger molecules than what is currently possible.

Our method does not require any prior knowledge of the payload’s structure. However, it does assume that each read contains a complete oligonucleotide, including its start and end primers. These primers are essential to initiate and terminate path-finding in the de Bruijn graphs. Unfortunately, some Illumina paired-end sequencing protocols do not retain this information. The base calling step typically removes primers, as they are biologically irrelevant in that context. Nevertheless, the oligonucleotide can be reconstructed by merging read pairs and appending artificial primers. As such, the ConClud software is compatible with both long- and short-read sequencing technologies.

DNA-based data storage is a promising solution for long-term, high-density information archiving. In this context, recovering high-quality sequences after the sequencing step is of utmost importance, as it directly influences the design of error-correcting codes tailored for DNA storage. When sequence reconstruction is highly accurate, the level of redundancy required for error correction can be significantly reduced, leading to substantial cost savings in the synthesis process. Moreover, being able to efficiently process large volumes of sequencing data is essential to ensure a smooth and scalable end-to-end data storage pipeline. Our work contributes to addressing both of these challenges, offering a reliable and efficient solution for DNA data storage systems.

## Acknowledgments

The work described in this paper is funded by the French ANR program ANR-22-PEXM-0005 MoleculArxiv and by the Labex CominLabs (dnarXiv project).

## Annex 1: ConCluD Software

Two versions of ConCluD exist: one designed for conventional multicore servers and one adapted to Processing-in-Memory architectures. Here, only the multicore version is considered. The source code is available here:

https://gitlab.inria.fr/pim/org.pim.dnarxiv/-/tree/main

## Annex 2: Generation of the IM1, IM10 and IM100 synthetic datasets

The datasets used for performance evaluation have been constructed from datasets archived in the following Zenodo repository:

https://zenodo.org/records/15387164

6 datasets that represents DNA encoded JPEG images are available:

1. **IM1-140-2**: 1 pool of 31,294 oligonucleotides of size 140 nt. All oligonucleotides share the same pair of primers (2 primers).
2. **IM1-248-2**: 1 pool of 35,090 oligonucleotides of size 248 nt. All oligonucleotides share the same pair of primers (2 primers).
3. **IM10-140-20**: 10 pools of 36,583 oligonucleotides of size 140 nt. Each pool have a different pair of primers (20 primers).
4. **IM10-248-20**: 10 pools of 35,069 oligonucleotides of size 248 nt. Each pool have a different pair of primers (20 primers).
5. **IM10-248-11**: 10 pools of 35,069 oligonucleotides of size 248 nt. Each pool have the same start primer but a different end primer (11 primers).
6. **IM100-248-20**: 100 pools of 35,114 oligonucleotides of size 248 nt. Each pool have a different pair of primers among 10 start primers and 10 end primers (20 primers).

The structure of an oligonucleotide is as follows:

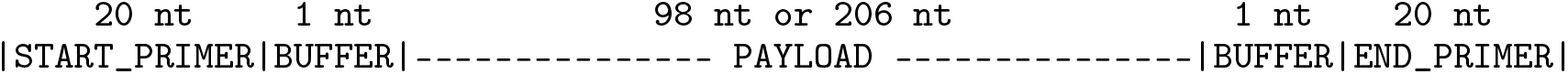

The length of the primers are 20 nt.

The length of the payload is 98 nt for oligos of size 140 nt and 206 nt for oligos of size 148 nt.

A 1-bit buffer is inserted between the payload and the primers due to biotechnological constraints. The payload contains encoded indexing and CRC information.

The scripts for generating the oligonucleotide datasets are available here:

https://gitlab.inria.fr/pim/org.pim.dnarxiv/-/tree/main/paper_scripts/datasets/IM-1-10-100

The generation of N pools of oligonucleotides from a binary file is performed as follows:

1. Split the binary file into N binary strings. N is determined by the target length of the oligonucleotides and the number of oligonucleotides by pool.
2. Apply a SHA256 hash to each binary strings with known key in order to obtain (pseudo) random binary strings of identical size, which is added to the binary string with an XOR operation. This operation ensures a good GC percentage balance and avoids large homopolymers.
3. Cut the binary strings into P fragments, relatively to the length of the oligonucleotides
4. Append a 16-bit index and 20-bit checksum to each fragment.
5. Convert fragments to DNA sequences with an encoding algorithm at a density of 1.66 bit/nt, ensuring a good GC balance and no homopolymeres.
6. Compute primers that do not hybridize with the fragments of a given pool.
7. Add primers and buffer to each fragment.

In order to be compliant with the DBGPS software, oligonucleotides from these datasets are modified. A 4 bytes index (= 16 bases) is inserted between the start primer and the payload, and a 2 bytes CRC (= 8 bases) is inserted between the payload and the end primer. The index is chosen to not have any homopolymers larger than 4 bases. The CRC is computed with the code of the public Python implementation of DBGPS. The helper scripts to convert the reference prints in the console the length and index start/end parameters to use to run ConCluD and DBGPS.

Long reads are generated with the PBSIM3 simulator, and short reads by the ART simulator. As a result we get the following synthetic datasets:

- IM1-164ont2 made from IM1-140-2 and PBSIM3
- IM1-164ill2 made from IM1-140-2 and ART
- IM1-272ont2 made from IM1-248-2 and PBSIM3
- IM10-164ont20 made from IM10-140-20 and PBSIM3
- IM10-272ont20 made from IM10-248-20 and PBSIM3
- IM10-272ont11 made from IM10-248-11 and PBSIM3
- IM100-272ont20 made from IM100-248-20 and PBSIM3

## Annex 3: Experiment Information

All the conditions and the implementation of the various experiments carried out within the framework of this article are available here:

https://gitlab.inria.fr/pim/org.pim.dnarxiv/-/tree/main/paper_scripts

